# Long-Term Effects of Radiation Therapy on Cerebral Microvessel Proteome: A Six-Month Post-Exposure Analysis

**DOI:** 10.1101/2025.01.13.632491

**Authors:** Vikram Subramanian, Denise Juhr, Piero Giansanti, Isabella M. Grumbach

## Abstract

**Background:** Radiation therapy (RT) treats primary and metastatic brain tumors, with about one million Americans surviving beyond six months post-treatment. However, up to 90% of survivors experience RT-induced cognitive impairment. Emerging evidence links cognitive decline to RT-induced endothelial dysfunction in brain microvessels, yet *in vivo* studies of endothelial injury remain limited. Investigating the molecular and cellular pathways connecting RT, endothelial dysfunction, and cognitive impairment is vital for developing targeted interventions. This study examines proteomic changes in cerebral microvessels following RT.

**Methods:** We conducted a comprehensive quantitative analysis comparing the proteome in cerebral microvessels from five control mice and five irradiated mice (12 Gy) 6 months after RT. Bioinformatics analyses included gene ontology (GO) enrichment, Mitocarta analysis, Ingenuity Pathway Analysis (IPA), and iPathwayGuide. Predictions from the analyses were validated by western blotting.

**Results:** Our data identified significant dysregulation of 414 proteins following RT, with 157 upregulated and 257 downregulated. Gene ontology analysis indicated that the majority of the dysregulated proteins were part of various metabolic pathways. Cross referencing with Mitocarta revealed a significant presence of mitochondrial proteins among the dysregulated proteins, indicating potential mitochondrial metabolic dysfunction. Further investigation with IPA analysis uncovered 76 enriched canonical pathways, 34 transcription regulators, 6 nuclear receptors, and 5 growth factors involved in RT-induced damage responses in cerebral microvessels. IPA canonical pathway analysis predicted mitochondrial dysfunction due to inhibition of various metabolic pathways in the irradiated group. Validation with western blotting confirmed the bioinformatics predictions from the proteomic dataset.

**Conclusions:** Our data show significant proteomic changes in cerebral microvessels 6 months post-radiation, including oxidative phosphorylation, the TCA cycle, and glycolysis, suggesting metabolic mechanisms of RT-induced microvascular dysfunction.

## 1 INTRODUCTION

Microvascular injury is a hallmark of late radiation damage across various organs depending on the radiation field. Examples include renal dysfunction, skin fibrosis and heart failure ^1–5^. One area where the effects of radiation on normal tissue are particularly apparent is injury after treatment of brain tumors ^6–8^. In fact, up to 90% of patients experience cognitive function decline within three to six months after radiation ^9–15^. Vascular damage, neuroinflammation and neuronal injury have been proposed as mechanisms driving cognitive decline ^16^. Microvascular injury can compromise the blood-brain barrier (BBB) breakdown, leading to inflammatory cell infiltration and inflammation ^17^. It can also result in apoptosis and senescence in vascular wall cells, potentially causing vascular rarefaction, reduced perfusion, and ischemia ^18^. While there is consensus on a role of microvascular injury, the exact mechanisms and extent of microvascular injury’s role in cognitive decline remains poorly understood.

In this study, we aimed to uncover changes in the microvascular proteome following RT. We chose to conduct our study using 12-month-old mice, approximately equivalent to 60-year-old humans, which is close to the reported median age of patients treated with radiation therapy for brain tumors ^19^. Of note, there is age-dependent variation in brain radiation injury, with the highest pathology occurring in the very young and elderly ^16,20^. We also opted for a six-month follow-up to focus on chronic changes from radiation injury. Previous work, including our own, has suggested that various pathways, such as mitochondrial injury, altered vascular transport, and changes in cell adhesion and tight junctions, may play a role in the long-term effects of radiation ^21–25^. To gain an unbiased view on long-term effects of radiation exposure on the cerebral microvessel proteome, we conducted bioinformatics analyses using Ingenuity Pathway Analysis (IPA) and iPathwayGuide to identify significantly affected canonical pathways that may contribute to cerebral microvasculature dysfunction. Additionally, we performed immunoblotting to validate our proteomics findings and bioinformatics results.

## 2 MATERIALS AND METHODS

### 2.1 Animals Housing and Irradiation Procedure

One-year-old male C57BL/6J mice were obtained from the Jackson Laboratory. The mice were randomly assigned to the treatment (RT) or the control group, each with five animals. They were housed in temperature-controlled rooms with a 12-hour light/dark cycle, provided standard rodent chow, and had water access *ad libitum*. To avoid gender-related confounding factors, only male mice were used. The control group mice (n=5) underwent sham irradiation, anesthetized with ketamine and xylazine, placed in the radiation chamber without exposure. Mice were randomly selected for irradiation versus sham treatment. The irradiated group (n=5) received a 12 Gy X-ray dose to the whole brain using the XStrahl Small Animal Radiation Research Platform, which uses a 60 kVp beam of 0.2 mm Al quality for Cone Beam CT and a 220 kVp 0.63 mm Cu quality beam for treatment. The 12 Gy dose corresponds to 2 Gy fractions of a 40 Gy total, commonly used for RT in patients with five or more brain metastasis. Both groups were euthanized six months post-irradiation for analysis. No mice were excluded from the analysis. All experimental procedures were approved by the Institutional Animal Care and Use Committees of the University of Iowa and the Iowa City VA Health Care System, following Institute for Laboratory Animal Research standards.

### 2.2 Isolation of Cerebral Microvessels and Preparation of Protein Samples

Mice were euthanized using 100% CO2 inhalation, followed by cervical dislocation. Brains from both control and irradiated groups were surgically removed, rinsed with cold Dulbecco’s phosphate-buffered saline (DPBS) (Gibco #2430024), snap-frozen in liquid nitrogen, and stored at −80 °C. Cerebral microvessels were isolated following Lee et al ^26^. Briefly, brain tissue was homogenized with a loose-fit Dounce grinder (Sigma-Aldrich #D9063) in MCDB 131 medium (Thermo Fisher Scientific #10372019) and centrifuged at 2,000 g for 15 minutes at 4 °C. The pellet was resuspended in a 15% (wt./vol) 70- kDa dextran solution (Sigma-Aldrich #31390) and centrifuged at 7,000 g for 15 minutes at 4 °C. The top layer containing myelin and parenchymal cells was discarded. Microvessels were collected using a 40-µm cell strainer (Corning #352340), rinsed with cold DPBS, and transferred into MCDB 131 medium containing 0.5% (wt./vol) BSA (Millipore Sigma #126609). The suspension was centrifuged at 16,100 g for 30 minutes at 4 °C to pellet the microvessels. Pellets were stored at −80 °C for analysis.

For protein extraction, the isolated microvessel pellets were mixed with 130 µl of RIPA buffer (Fisher Scientific #R0278) containing protease (Pierce™ #A32963) and phosphatase inhibitor cocktails (Pierce™ #A32957). The mixture was vortexed three times for 30 seconds each (Fisher Scientific #0215370), then agitated for 30 minutes at 15-20 °C. Samples were vortexed again and heated at 95 °C for 10 minutes. To further break down the tissue, the samples were lysed using a Covaris E220 focused ultrasonicator (Covaris #500239) with following parameters. The instrument parameters for shearing were as follows: water level set point 10, water temperature 6°C, peak incident power 175 W, duty factor 10%, cycles per burst 200, and duration 300 s. Homogenized lysates were transferred to Eppendorf tubes and centrifuged at 16,100 g (Eppendorf #5415R) for 30 minutes. The supernatant was collected and stored at −80 °C. Total protein concentration was measured using a BCA protein assay kit (Pierce™ #23225). Finally, 30 µg of protein from each sample was used for quantitative proteomics analysis. Samples were labeled with a numeric identifier that did not include information about irradiation or sham treatment and sent for proteomic analysis.

### 2.3 Protein Digestion

Protein digestion was performed using the SP3 method ^27^. For each sample, 30 μg of protein in 150 μL of lysis buffer was incubated with 10 μL of SP3 beads (a 1:1 mixture of Sera-Mag Speed Beads A and B, Thermo Scientific). Pure acetonitrile (ACN, VWR Chemicals) was added to a final concentration of 70% (v/v). The samples were incubated mixed at 800 rpm for 18 minutes, then placed on a magnet rack for 3 minutes to immobilize the SP3 beads. The supernatant was discarded, and the SP3 beads were washed three times with 1 mL of 80% ethanol/water (v/v) and once with 800 μL of ACN. Bound proteins were reduced with 100 μL of 10 mM 1,4-dithiothreitol (DTT, Sigma) in 50 mM ammonium bicarbonate (Sigma), pH 8.0, and incubated at 37 °C with shaking at 800 rpm for 1 hour. Proteins were then alkylated with 55 mM 2-chloroacetamide (CAA, Sigma) for 1 hour at 37 °C in the dark. Finally, 1 μg of trypsin (Thermo Scientific) was added to each sample, and they were incubated overnight at 37 °C with shaking at 800 rpm. Digests were acidified with formic acid (FA, Carlo Erba) to a final concentration of 1% (v/v), dried in vacuo, and stored at −80 °C until use.

### 2.4 Automated Off-line Fractionation and LC-MS/MS Analysis

Peptides were re-suspended in 110 μL of buffer A (25 mM ammonium formate, pH 10) and subjected to high pH reverse-phase fractionation using the AssayMAP Bravo platform and 5 mL RP-S cartridges (Agilent). Cartridges were primed with 150 μL each of isopropanol, acetonitrile (ACN), and buffer B (80% ACN in 10 mM ammonium formate, pH 10) at a 50 μL/min flow rate. They were then equilibrated with 100 μL of buffer A, and peptides were loaded at 5 μL/min, collecting the flow-through (FT). Peptides were eluted with 25 mM ammonium formate, pH 10, using increasing ACN concentrations (5%, 10%, 15%, 20%, 25%, 30%, 80%). The seven fractions were pooled into four final fractions, dried in vacuo, and stored at −80 °C.

Nano-flow LC-MS/MS was performed using a Dionex Ultimate 3000 UHPLC+ system coupled to an Orbitrap Eclipse mass spectrometer (Thermo Fisher Scientific). Peptides were first delivered to a trap column (75 μm i.d. × 2 cm, packed in-house with 5 μm Reprosil C18 beads) and washed with 0.1% formic acid (FA) at 5 μL/min for 10 minutes, then transferred to an analytical column (75 μm i.d. × 45 cm, packed in-house with 3 μm Reprosil C18 beads) at 300 nL/min. Chromatographic separation used a linear gradient of solvent B (0.1% FA, 5% dimethyl sulfoxide (DMSO) in ACN) and solvent A (0.1% FA, 5% DMSO in water) over a 90-minute total run time.

Full-scan MS spectra were recorded in the Orbitrap from 360 to 1,300 m/z at 60,000 resolutions, with an automatic gain control (AGC) target of 100% and a maximum injection time (maxIT) of 50 ms. The most intense precursors were isolated with a 1.3 m/z isolation window for high-energy collisional dissociation (HCD) fragmentation. Fragment ions were recorded in the Orbitrap at 15,000 resolutions, with a maxIT of 22 ms and an AGC target of 200%. Normalized collision energy (NCE) was set to 25%.

Charge state screening selected precursors with charge states 2 to 6 for fragmentation, within a 2- second cycle time. Dynamic exclusion was set to 35 seconds.

### 2.5 Identification and Quantitation of Peptides and Proteins

Raw mass spectrometry data were processed using MaxQuant (version 2.2.0.0) with its built-in search engine, Andromeda ^28^. Spectra were searched against the UniProtKB database for Mus musculus (UP000000589, 55,338 entries, downloaded October 2022). Enzyme specificity was set to trypsin, allowing up to two missed cleavages. The search included cysteine carbamidomethylation as a fixed modification, and protein N-term acetylation and methionine oxidation as variable modifications. Identifications were filtered to achieve a 1% false discovery rate (FDR) at protein and peptide levels. The "match between runs" and "second peptide" options were enabled. Label-Free Quantification (LFQ) was performed using the MaxLFQ algorithm ^29^. The mass spectrometry proteomics data have been deposited in the ProteomeXchange Consortium via the PRIDE repository ^30^ with the dataset identifier PXD058732 (Website: http://www.ebi.ac.uk/pride, Username, Password: RJJaEf7seviy).

### 2.6 Proteomics Data Analysis and Pathway Enrichment Analysis

Data analysis was performed using Maxquant and Perseus software (version 2.0.7.0). Protein identifications were filtered to remove contaminants and decoy hits before normalizing log2-transformed LFQ intensity values by median centering (Supplemental Table S1). Only proteins quantified in all biological replicates were retained for statistical analysis **(n = 2,457, Supplemental Table S2).** Significantly dysregulated proteins were identified using a Student’s t-test with an S0 parameter of 0.1, with p-values adjusted using a permutation-based FDR of 5%. For the final quantifications, proteins identified by at least two unique peptides and demonstrating a fold change ratio of <0.77 or >1.30, with corrected p-values according to prior studies were defined as being significantly differentially expressed and used for all bioinformatics analysis **(n = 414, Supplemental Table S3)** ^31–33^. Hierarchical cluster analysis of significantly dysregulated proteins was performed using SR plot ^34^. A volcano plot illustrating differentially expressed proteins was generated using VolcaNoseR ^35^. Enrichment analysis of dysregulated proteins was conducted to assess their impact on biological processes, cellular locations, molecular functions, and KEGG pathways after radiation exposure, using HemI 2.0 ^36^. Canonical pathway, upstream regulator and disease and biofunction analyses were performed with INGENUITY Pathway Analysis (IPA) software ^37^, applying a threshold of –log (p-value) >1.3. Pathways with a z-score >2.0 were considered activated, while those with a z-score <–2.0 were inhibited. Impact pathway analysis on differentially expressed proteins was conducted using iPathwayGuide ^38^ with pathway annotations from the KEGG database and gene ontology annotations from the Gene Ontology Consortium database. A threshold of FDR-corrected p-value <0.05 was used to identify significantly affected pathways.

### 2.7 Immunoblotting Analysis

Equal amounts of protein (10 µg) from control and irradiated groups were separated on NuPAGE 4% to 12% Bis-Tris gels (Life Technologies) and transferred to polyvinylidene difluoride (PVDF) membranes using the Mini-PROTEAN 3 blotting system (Bio-Rad). Membranes were stained with Ponceau S (Cell Signaling #59803S) to confirm protein transfer, washed in TBS with Tween™ (TBST), and blocked with 3% BSA (RPI Research #9048-46-8) or Starting Block™ Blocking Buffer (Thermo Fisher Scientific #37538) for 2 hours. Membranes were incubated overnight at 4°C with primary antibodies, including anti-NDUFB8 (Abcam, no. ab110413), anti-NDUFA11 (ABclonal, no. A16239), anti-NDUFA4 (ABclonal, no. A15693), anti-SDHA (Cell Signaling, no. 11998), anti-SDHB (Abcam, no. ab110413), anti-UQCRC2 (Aviva systems biology, no. OAAN01132), anti-MT-CO2 (ABclonal, no. A11154), anti-COX6B1 (ABclonal, no. A2228), anti-ATP5F1C (ABclonal, no. A15257), anti-ATP6VID (ABclonal, no. A12940), anti-ATP5F1A (ABclonal, no. A11217), anti-ACLY (ABclonal, no. A3719), anti-IDH3A (ABclonal, no. A14650), anti-DLD (ABclonal, no. A13296), anti-ACO2 (ABclonal, no. A3716), anti-MDH2 (ABclonal, no. A13516), anti-ENO2 (Cell Signaling, no. 9536), anti-TPI (ABclonal, no. A15733), anti-MPC1 (Cell Signaling, no.14462), and anti-MPC2 (Cell Signaling, no. 46141). After washing, membranes were incubated with secondary antibodies at a 1:5000 dilution for 2 hours at room temperature. After washing, membranes were developed using the Cytiva Amersham™ ECL™ Prime Western Blotting Detection Reagent and scanned. Immunoblot bands from five biological replicates in both control and irradiated groups were analyzed and quantified using ImageJ software (http://rsbweb.nih.gov/ij/).

### 2.9 Statistical Analysis

All experiments were conducted in five biological replicates. Data were presented as mean with standard error, and statistical analyses were conducted using GraphPad Prism 9.0 software. Normal distribution was evaluated using the D’Agostino-Pearson omnibus normality test. Statistical significance for comparisons between two groups was determined using the non-parametric Mann-Whitney test. Differences were considered significant if p-values were <0.05.

## 3 RESULTS

### 3.1 Proteome profiling of cerebral microvessel after radiation therapy

To investigate the effects of radiation on cerebral microvasculature, we performed quantitative proteomics using LC-MS/MS on samples from mice six months after RT with 12 Gy x-ray or sham treatment **(Figure 1A)**. The full workflow for this proteomics study is shown in **Figure 1B**. A total of 7,462 proteins were detected, and 2,457 were quantified **(see Supplementary Table S1, S2)**. Label-free quantitation (LFQ) demonstrated reproducibility among biological replicates from both control and irradiated mice, as shown by Pearson correlation **(Figure 1C)**. Principal component analysis (PCA) using LFQ intensity values of all identified proteins revealed a clear separation between control and irradiated groups, indicating significant changes in protein expression due to radiation **(Figure 1D)**.

**Figure 1:**
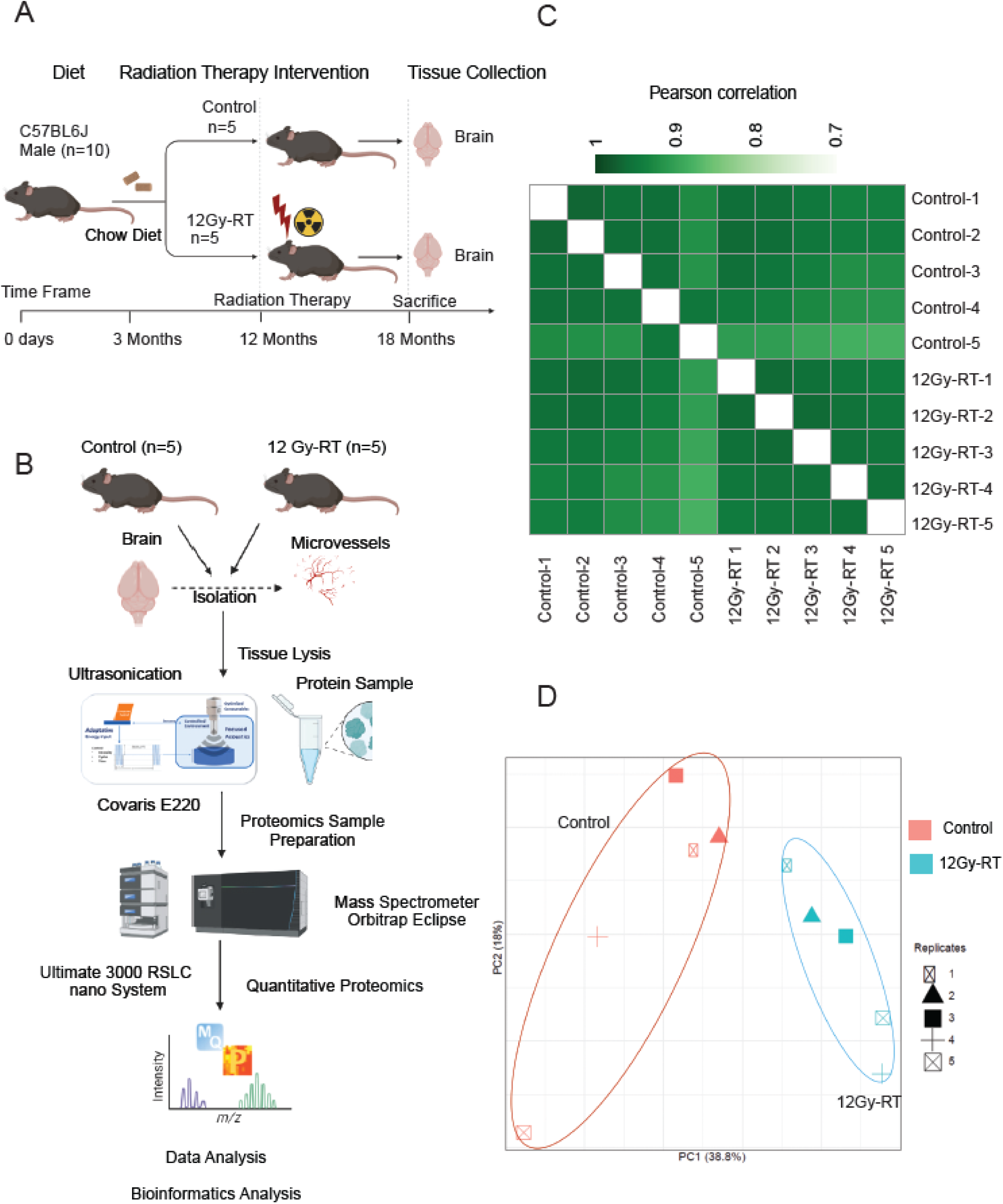
Quantitative proteomics analysis of the cerebral microvessels. **(A)** Workflow for radiation therapy (RT) and sample collection. **(B)** Workflow for high-throughput identification and quantification of proteins from cerebral microvessels in control and irradiated mice (12 Gy-RT). **(C)** Pearson correlation matrix based on label-free quantification (LFQ) intensities, showing sample reproducibility in control and irradiated groups. **(D)** Principal component analysis (PCA) plot for proteins identified in biological replicates of control and irradiated groups.

Proteins were classified as identified with at least one valid MS/MS spectrum, quantified if detected in all five biological replicates, and dysregulated if their relative abundance significantly differed between the groups. Among the 2,457 quantified proteins, 414 proteins were significantly dysregulated between irradiated and control groups, and these proteins were used for further bioinformatics analysis. Hierarchical clustering of z-scored LFQ intensities of dysregulated proteins highlighted substantial expression differences between the groups **(Figure 2A)**. Of the 414 dysregulated proteins, 157 were upregulated, and 257 were downregulated **(see Supplementary Table S3)**. Volcano plots depicting these changes, with the top 10 proteins in each category labeled, are shown in **Figure 2B**.

**Figure 2:**
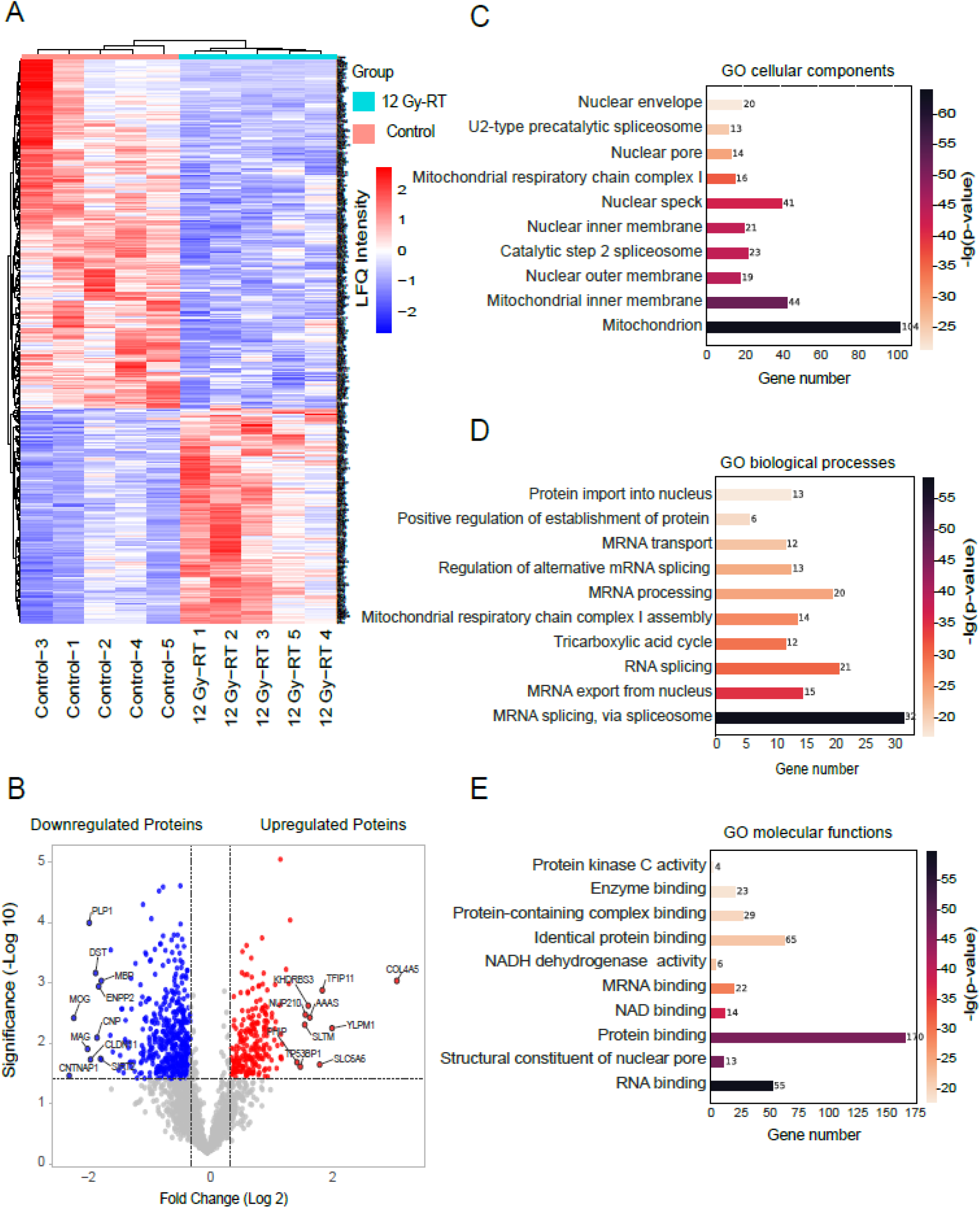
Significant differences in the cerebral microvessel proteomes between control and irradiated groups. **(A)** Hierarchical clustering analysis (heatmap) using unsupervised Euclidean distance for differentially expressed proteins across biological replicates. Z-scored LFQ intensities for individual mice are shown side by side. **(B)** Volcano plots of differentially expressed proteins, with the y-axis representing −log10 p-value significance and the x-axis representing log2 fold change. The top 10 proteins with the most significant decrease and increase in expression are highlighted in blue and red, respectively (p < 0.05). The dotted line indicates the cut-off for protein expression fold change (log2) against statistical significance [-log10 (p-value)]. **(C)** GO cellular components, **(D)** GO biological processes, and **(E)** GO molecular function analysis of significantly dysregulated proteins compared to the control group. The top 10 enriched GO terms are displayed with corrected p-values and gene numbers. The colored bar indicates enriched terms with corresponding corrected p-values and gene numbers. Analyses were performed using the HemI 2.0 web tool.

To investigate the biological significance of the dysregulated proteins, we utilized the Helm webtool for further analysis. The highest representation of dysregulated proteins was observed in the cellular components “mitochondrion”, “mitochondrial inner membrane”, and “nuclear speck”, a specialized subnuclear structure associated with splicing factors (**Figure 2C**). Regarding biological processes, proteins involved in mRNA splicing and processing, as well as the TCA cycle and mitochondrial respiratory chain complex I assembly, were predominantly affected (**Figure 2D**). For molecular function, dysregulated proteins were primarily associated with protein and RNA binding activities (**Figure 2E**). A complete list of these categories and their corrected p-values is provided in Supplementary Table S4.

### 3.2 Impact Pathway Analysis

In addition, we analyzed all changes with iPathwayGuide that unlike other functional analysis tools, considers a gene’s position within each pathway and its interactions with other genes. The analysis revealed that the significantly dysregulated proteins were enriched in 266 pathways, with strongest impact on metabolic pathways, nucleocytoplasmic transport, one carbon metabolism, oxidative phosphorylation, spliceosome, and the tricarboxylic acid cycle **(Figure 3A-F)**. Additionally, several non-metabolic pathways were significantly enriched in irradiated cerebral microvessels, including cGMP-PKG signaling, vascular smooth muscle contraction, mRNA surveillance, calcium signaling **(Supplemental Figure 1A-E)**, focal adhesion, tight junction, and gap junction, **(Supplemental Figure 2A-C)**. Moreover, proteins enriched in the spliceosome and mRNA surveillance pathways were upregulated in irradiated group, potentially suggesting an increased stress response and errors in mRNA processing due to radiation exposure. A complete list of enriched pathways and their corrected p-values is available in **Supplementary Table S5**.

**Figure 3:**
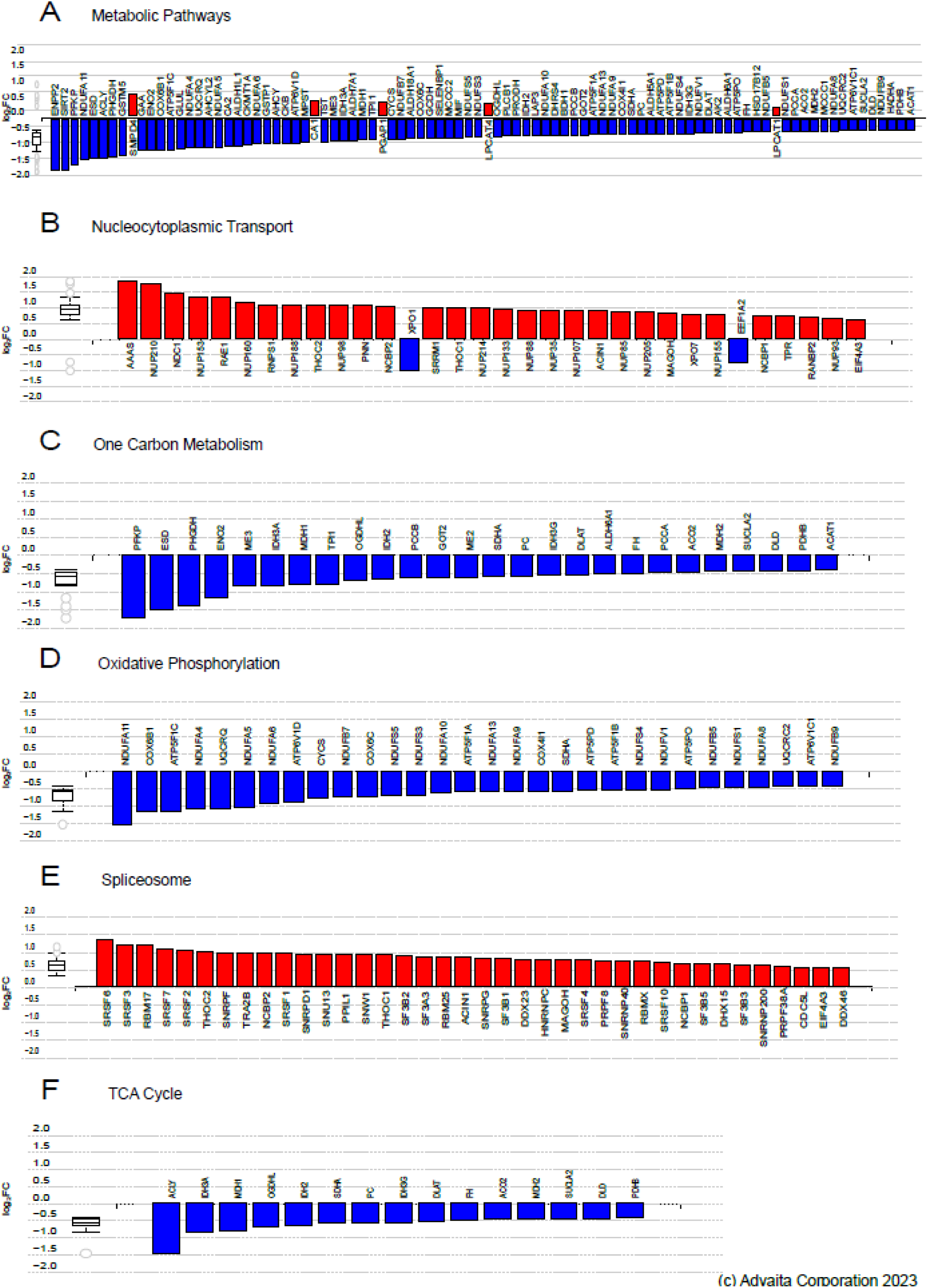
Impact pathway (iPathway guide) analysis of dysregulated proteins by RT. **(A-F)** Bar graphs showing dysregulated proteins mapped to pathways identified by iPathway analysis based on FDR-corrected p-value significance: **(A)** Metabolic pathways, **(B)** nucleocytoplasmic transport, **(C)** one carbon metabolism, **(D)** oxidative phosphorylation**, (E)** spliceosome, and **(F)** TCA cycle. Genes are ranked by absolute log-fold change, with upregulated genes in red and downregulated genes in blue. Box-and-whisker plots summarize the distribution of differentially expressed genes in each pathway, with boxes representing the 1st quartile, median, and 3rd quartile, and circles indicating outliers.

The core analysis module in the Ingenuity Pathway Analysis (IPA) software was used to identify altered canonical pathways in cerebral microvessels following radiation therapy (RT). Using a -log (p-value) cutoff of 1.3 (Fisher’s exact test) and a z-score threshold (Z-score > 2 for activation, Z-score < -2 for inhibition), 76 enriched canonical pathways were identified. Of these, 68 pathways were predicted to be inhibited and 8 activated post-RT. **Figure 4A-C** and **Supplementary Table S6** highlight 15 canonical pathways associated with cerebral microvessel function. Based on altered protein expression, the three most significantly inhibited pathways were oxidative phosphorylation, the TCA cycle, and Gαq signaling, while the five most significantly activated pathways included mitochondrial dysfunction, sirtuin signaling, granzyme A signaling, the spliceosomal cycle, and RHOGDI signaling.

**Figure 4:**
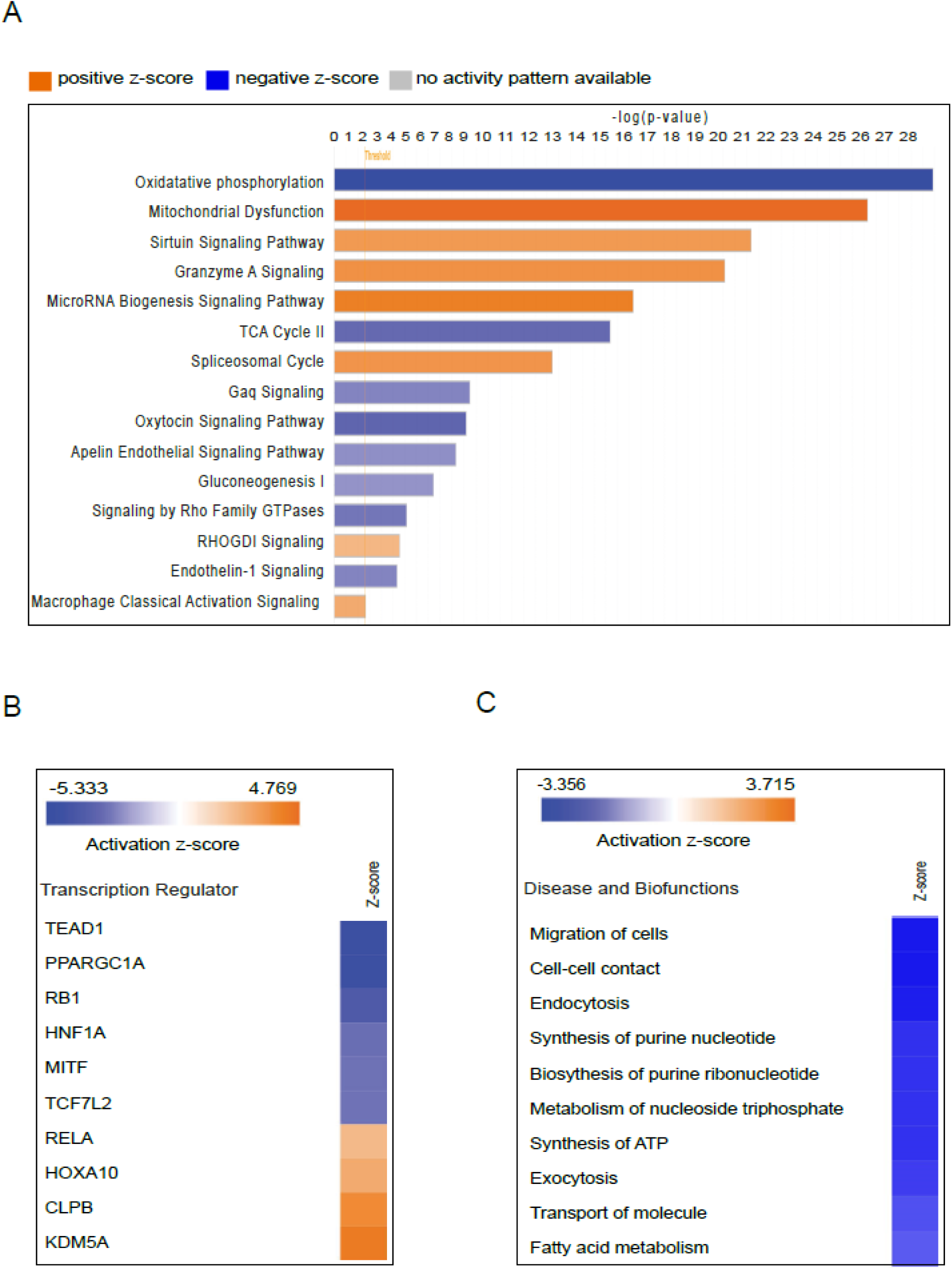
Ingenuity pathway analysis (IPA) of all dysregulated proteins from the irradiated group. **(A)** Fifteen selected canonical pathways predicted to be activated or inhibited based on z-scores and a B–H p-value < 0.05 (calculated using Fisher’s exact test and adjusted with the Benjamini-Hochberg method). A positive z-score (orange) indicates pathway activation, while a negative z-score (blue) indicates inhibition. Longer bars indicate stronger significance (source: http://www.INGENUITY.com). **(B)** Heatmap of the top 10 transcription regulators predicted to be activated (z-scores > 2) or inhibited (z-scores < -2). **(C)** Predicted activation or inhibition of selected diseases and biofunctions based on z-scores (> 2 for activation; < -2 for inhibition) in the irradiated group.

An upstream regulator analysis using the z-score algorithm identified 34 enriched transcription regulators in irradiated cerebral microvessels. The top activated regulators were KDM5A (Z-score = 4.243), CLPB (Z-score = 3.742), HOXA10 (Z-score = 2.714), and RELA (Z-score = 2.201), while the top inhibited regulators included TEAD1 (Z-score = -4.899), PPARGC1A (Z-score = -4.872), RB1 (Z-score = -3.90), HNF1A (Z-score = -2.985), and MITF (Z-score = -2.887) (**Figure 4B**).

A disease and biofunction analysis of 414 differentially expressed proteins revealed enrichment for 41 diseases and biological functions post-radiation exposure. Of these, 31 functions were predicted to be inhibited, while 10 were activated. **Figure 4C** depicts the selected 10 enriched diseases/biological functions, associated molecules, and their p-values and z-scores. Notable affected functions in cerebral microvessels after radiation therapy included cell migration, cell-to-cell contact, ATP synthesis, molecule transport, and fatty acid metabolism, as detailed in **Supplementary Table S6**.

### 3.3 RT induces changes in the expression of mitochondrial proteins within Cerebral Microvessels

Given the strong evidence for changes in the mitochondrial proteome by different analysis tools, we investigated the impact of RT on mitochondrial protein expression and associated functions in more detail. We mapped all 2,467 quantified cerebral microvessel proteins to the Mouse Mitocarta3.0 database. This analysis identified 385 mitochondrial proteins, with 84 significantly dysregulated (all downregulated) in the irradiated group compared to controls **(Figure 5A; Supplementary Table S7)**.

**Figure 5:**
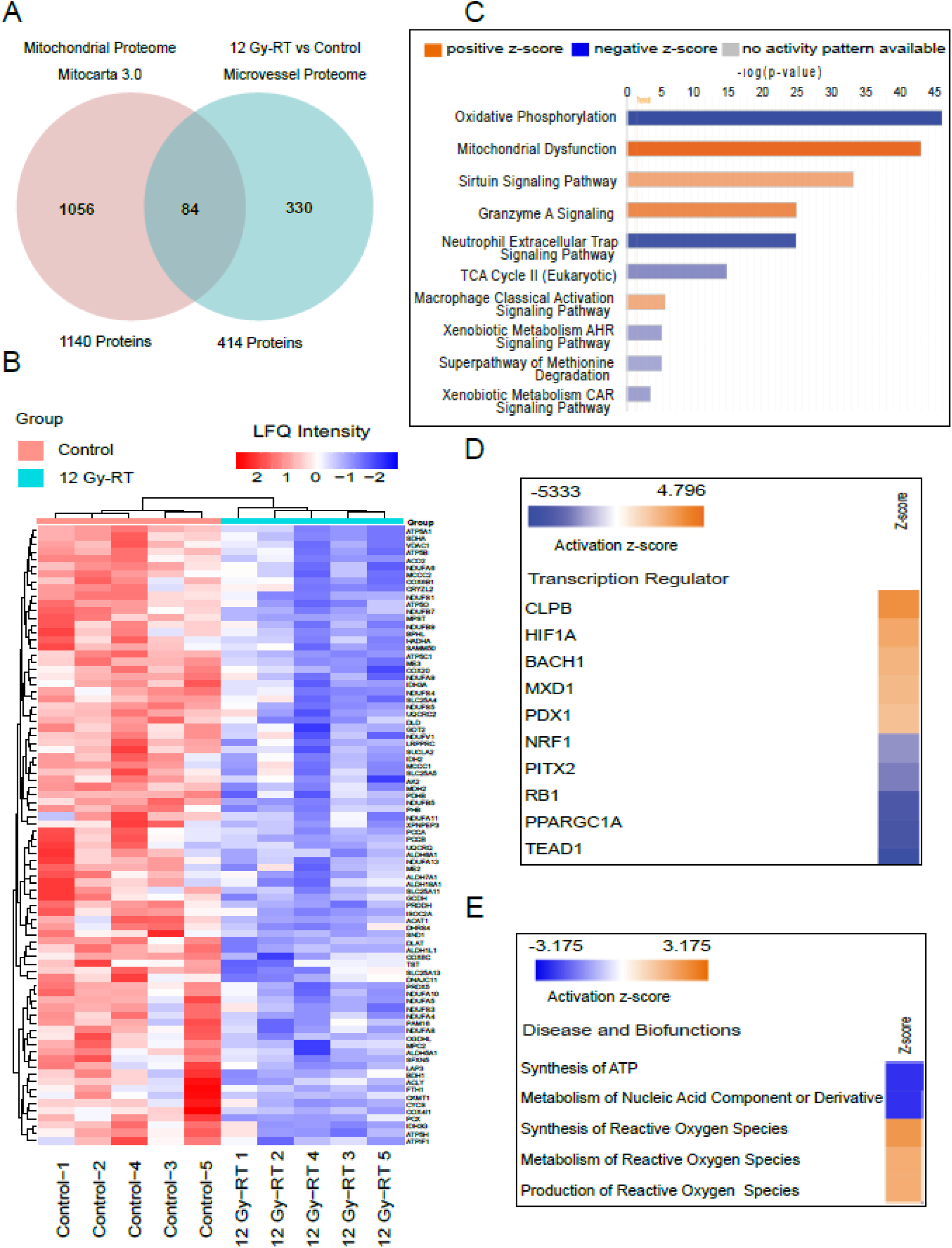
Ingenuity pathway analysis (IPA) of mitochondrial proteins that are dysregulated in the irradiated group. **(A)** Venn diagram showing 84 mitochondrial proteins shared between the mitochondrial proteome database (Mitocarta 3.0) and the cerebral microvessel proteome. **(B)** Hierarchical clustering analysis (heatmap) of differentially expressed mitochondrial proteins across biological replicates from control and irradiated groups, with z-scored LFQ intensities for individual mice. **(C)** Top 10 predicted canonical pathways based on z-scores and B–H p-values < 0.05 (calculated using Fisher’s exact test and adjusted with the Benjamini-Hochberg method). A positive z-score (orange) indicates pathway activation, while a negative z-score (blue) indicates inhibition (source: http://www.INGENUITY.com). **(D)** Heatmap of the top 10 transcription regulators predicted to be activated (z-scores > 2) or inhibited (z-scores < -2). **(E)** Predicted activation or inhibition of selected diseases and biofunctions based on z-scores (> 2 for activation; < -2 for inhibition) in the irradiated group.

We created a hierarchical cluster heatmap using the LFQ intensity of significantly dysregulated mitochondrial proteins, clearly separating control and irradiated groups (**Figure 5B**). To confirm the biological significance of these changes and their impact on mitochondrial function, we performed gene ontology (GO) analysis with the Helm webtool. The most affected proteins were linked to the mitochondrial inner membrane, mitochondrial respiratory chain complex I, and mitochondrial matrix (**Supplementary Figure 3A, Supplementary Table S8**). Key impacted biological processes included mitochondrial respiration, the tricarboxylic acid cycle, and 2-oxoglutarate metabolic process (**Supplementary Figure 3B**).

Using IPA core analysis, we examined affected canonical pathways in the mitochondrial protein data set. This analysis identified 14 enriched pathways, 10 predicted to be inhibited and 4 activated following radiation treatment. The top three inhibited pathways were oxidative phosphorylation, neutrophil extracellular trap signaling, and the TCA cycle (**Figure 5C**). Additionally, upstream regulator analysis for transcription factors associated with mitochondrial proteins identified 14 enriched regulators. Among those, CLPB, and HIF1A were the top activated regulators, while TEAD1, PPARGC1A, and RB1 were the most inhibited (**Figure 5D**). Supplementary Table S9 provides a comprehensive list of canonical pathways, upstream regulators, and associated p-values.

Lastly, we used IPA software to map altered diseases and biological functions based on dysregulated mitochondrial proteins. The analysis highlighted 22 enriched diseases and functions, 17 predicted to be inhibited and 5 activated. Notably, ATP synthesis and nucleic acid metabolism were predicted to decrease, while synthesis, production and metabolism of reactive oxygen species were predicted to increase (**Figure 5E**).

### 3.4 Metabolic pathways are affected in cerebral microvessels after RT

Energy metabolism is essential for biosynthetic processes to facilitate the restoration of cell damage. While previous studies have extensively explored the impact of radiation on cardiac energy metabolism ^31,39,40^ and metabolic pathways in other tissues ^41,42^ the impact of radiation on cerebral microvessel metabolism is less understood. Our proteomic analysis revealed altered expression of proteins linked to key metabolic pathways, including oxidative phosphorylation (n=29), the TCA cycle (n=15), and glycolysis (n=7) (**Supplementary Figure 4A-C**). IPA analysis suggested inhibition of oxidative phosphorylation, the TCA cycle, glycolysis, and gluconeogenesis. To confirm these findings, we used western blotting to analyze dysregulated proteins in oxidative phosphorylation, the TCA cycle, and glycolysis.

Our analysis revealed radiation-induced changes in the expression of proteins across various OXPHOS system complexes. Western blot images for different OXPHOS protein subunits with adjustment for loading by Ponceau-stained whole membranes revealed significant downregulation of complex I proteins NDUFA11 and NDUFB8 (**Figure 6A-C**), complex II protein SDHB (**Figure 6A-C**), and complex IV protein MT-CO2 (**Figure 6A-C**) in irradiated samples. Although downregulation of complex III and V proteins ATP5F1C, ATP6VID, and ATP5F1A was observed, these changes did not reach statistical significance (**Figure 6A-C**).

**Figure 6:**
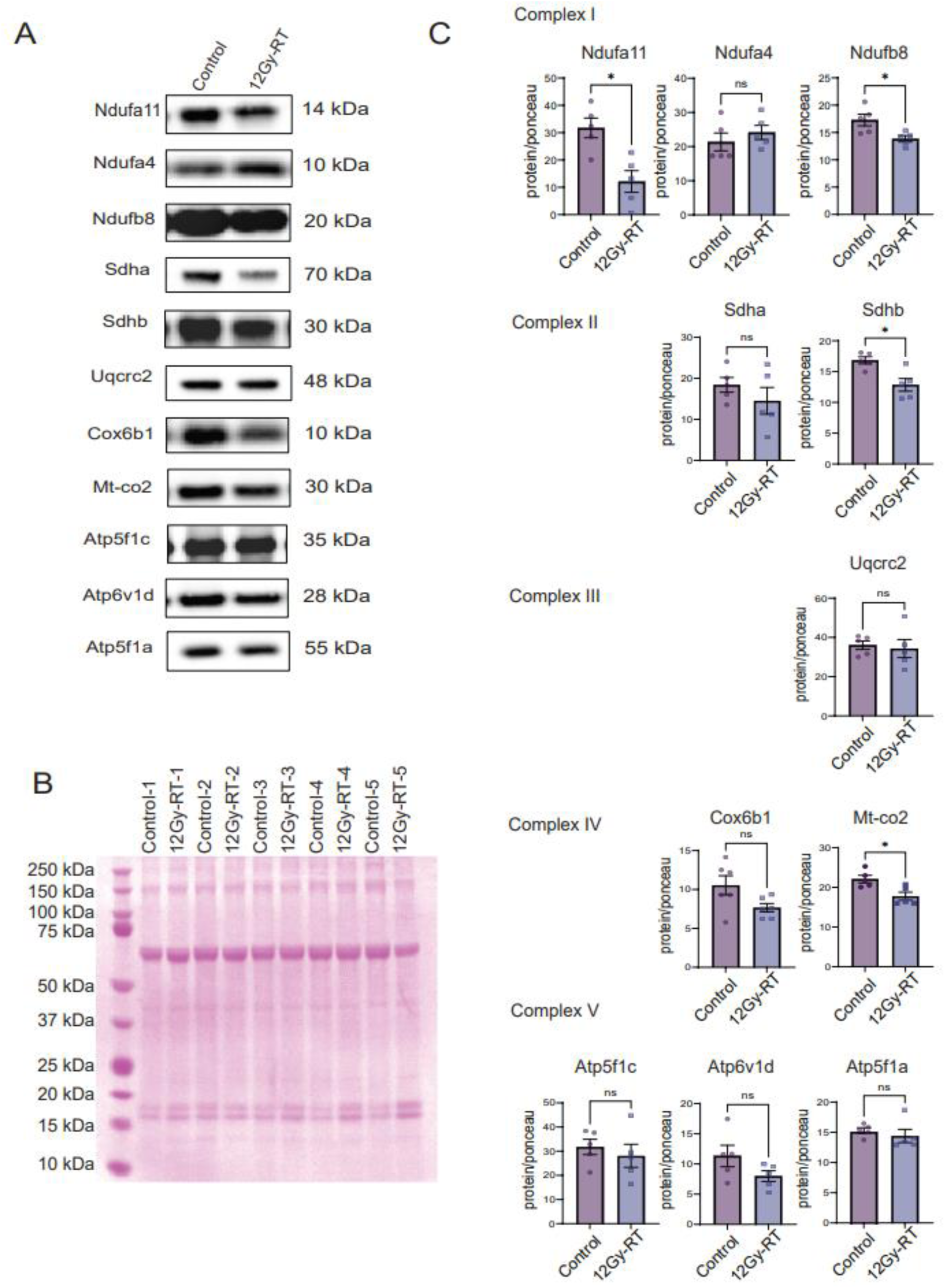
Validation of Metabolic Protein Expression Changes by Immunoblot. **(A)** Representative immunoblot images for OXPHOS complex subunits in control and irradiated groups. **(B)** Ponceau-stained membrane used as loading control. **(C)** Immunoblots of ETC complex I proteins (NDUFA11, NDUFA4, NDUFB8), complex II proteins (SDHA, SDHB), complex III protein (UQCRC2), complex IV proteins (COX6B1, MT-CO2), and complex V proteins (ATP5F1C, ATP6VID, ATP5F1A) in control and irradiated groups. Quantification of protein levels after normalization to Ponceau-stained membrane. Analysis by Mann-Whitney test.

Lastly, we examined proteins involved in the TCA cycle, glycolysis, and pyruvate transport (**Supplemental Figure 5A**) and detected non-significant trends towards decreases for TCA cycle proteins UQCRC2, ACLY, ACO2, IDH3A, DLD, SDHA, and MDH2 in the irradiated group (**Supplemental Figure 5B**). Similarly, analysis of glycolysis and pyruvate transport proteins revealed similar trends for glycolytic proteins TPI and ENOL2 as well as mitochondrial pyruvate carrier proteins MPC1 and MPC2 in the irradiated group (**Supplemental Figure 5C, 5D**).

## 4 DISCUSSION

Our study provides the first in-depth analysis of long-term proteome changes in non-tumor cerebral microvessels following RT in C57Bl6J mice, offering key insights into the impact of radiation on vascular health. A total of 414 dysregulated proteins were identified, with 257 showing downregulation. These findings highlight mitochondrial dysfunction and metabolic disruptions as central themes, with notable effects on pathways such as oxidative phosphorylation, the TCA cycle, and glycolysis.

Pathway enrichment analysis revealed significant alterations in 266 pathways. Among 76 canonical pathways, 68 were inhibited (e.g., oxidative phosphorylation), while 8 were activated (e.g., mitochondrial dysfunction). Mitochondrial proteins were particularly impacted, with a detailed analysis identifying 84 downregulated proteins linked to the mitochondrial inner membrane and respiratory chain complex I, indicating disrupted energy metabolism.

Proteomic changes also revealed inhibition of ATP synthesis, cell migration, and other cellular processes through pathways such as nucleo-cytosolic transport and spliceosomal activity. Increased oxidative stress was evidenced by the activation of reactive oxygen species synthesis, potentially as a result of mitochondrial dysfunction. Enriched disease-related processes further pointed to disruptions in molecule transport and fatty acid metabolism. Western blot analysis of OXPHOS proteins confirmed inhibited energy metabolism and mitochondrial dysfunction. These findings underscore the lasting effects of RT caused by RT in cerebral microvessels, particularly on mitochondrial health and metabolic pathways.

Given the strong impact of RT on the mitochondrial proteome, our study focused on changes in the mitochondrial and metabolic proteome. In most vascular beds, endothelial cells (ECs) predominantly use glycolysis and depend minimally on mitochondrial OXPHOS for ATP generation, as mitochondrial volume in ECs constitutes only 2–6% of total cellular volume ^43,44^. However, cerebromicrovascular ECs at the BBB possess nearly double the mitochondrial volume compared to other vascular beds ^45^. The BBB represents a selective yet dynamic interface between the blood and central nervous system that rigorously maintains neuronal homeostasis by regulating the transport of substances to and from the brain ^46,47^. The cerebral microvasculature engages in aerobic respiration to support the energy demands required for maintaining transport systems and barrier function ^48^. Doll and colleagues reported that mitochondrial “crisis” induced by a lipopolysaccharide challenge in cerebromicrovascular ECs significantly compromises BBB function in vitro and in vivo ^47^ but similar data after RT have been missing so far.

Since glycolysis is the primary metabolic pathway for energy production in ECs, it remains to be established how the extensive alterations in metabolic function following RT affect EC phenotypes. As such, further studies are needed to explore the positive link between energy depletion and EC dysfunction, for example, whether energy deficiency drives or results from long-term changes post-RT, including altered BBB permeability, cellular senescence, impaired angiogenesis, increased inflammation, reduced nitric oxide production, and a pro-thrombotic state ^43,49–53^. Since these changes may contribute to complications like neuroinflammation and ischemia in irradiated areas, largely due to the disruption of vascular homeostasis, targeting metabolic pathways may offer a novel approach to mitigating these long-term side effects.

Beyond metabolism, mitochondrial signaling is gaining recognition for its role in RT-induced tissue injury ^54–57^. While mitochondrial reactive oxygen species (ROS) have been highlighted as mediators of mitochondrial DNA damage and ETC dysfunction in ECs, additional factors, such as altered mitochondrial calcium in- and efflux, may also contribute ^22,56,58,59^. Moreover, mitochondrial DNA fragments as damage-associated molecular patterns may drive inflammatory signaling cascades after RT ^60^.

Additional to changes in the mitochondrial and metabolic proteome, our findings highlight dysregulation in proteins related to mRNA splicing and stress-response pathways, suggesting that RT drives transcriptional and post-transcriptional modifications ^61^. These changes potentially disrupt cellular homeostasis and protein synthesis, aggravating vascular dysfunction and warrant further investigation.

This study has several limitations. Firstly, exclusively male mice were used. Given merging evidence for differences in radiosensitivity in males versus females, findings should be validated in female mice ^62^. Secondly, the contributions of different cell types, such as pericytes, astrocytes, and ECs, cannot be fully resolved with the current methods. Thirdly, observations were limited to a single time point; future studies should examine various time points to provide a more comprehensive understanding of changes over time. Lastly, additional mechanistic studies are needed to confirm proteome study predictions about EC phenotypes, including respiration, barrier dysfunction, and mRNA processing.

In conclusion, our study provides evidence for mitochondrial dysfunction and dysregulation of protein synthesis pathways in cerebromicrovascular ECs following RT. These findings shed light on potential mechanisms underlying radiation-induced BBB breakdown and highlight the need for further investigation into the role of mitochondria in ECs. They also underscore the need for further research into the relationship between mitochondrial integrity, metabolic function, and vascular health in RT contexts.

